# In-cell processing enables rapid and in-depth proteome analysis of low-input *Caenorhabditis elegans*

**DOI:** 10.1101/2024.09.18.613705

**Authors:** Malek Elsayyid, Jessica E. Tanis, Yanbao Yu

**Affiliations:** Department of Biological Sciences, University of Delaware, Newark, DE 19716, USA; Department of Chemistry and Biochemistry, University of Delaware, Newark, DE, 19716, USA

## Abstract

*Caenorhabditis elegans* is a widely used genetic model organism, however, the worm cuticle complicates extraction of intracellular proteins, a prerequisite for typical bottom-up proteomics. Conventional physical disruption procedures are not only time-consuming, but can also cause significant sample loss, making it difficult to perform proteomics with low-input samples. Here, for the first time, we present an on-filter in-cell (OFIC) processing approach, which can digest *C. elegans* proteins directly in the cells of the organism after methanol fixation. With OFIC processing and single-shot LCMS analysis, we identified over 9,400 proteins from a sample of only 200 worms, the largest *C. elegans* proteome reported to date that did not require fractionation or enrichment. We systematically evaluated the performance of the OFIC approach by comparing it with conventional lysis-based methods. Our data suggest equivalent and unbiased performance of OFIC processing for *C. elegans* proteome identification and quantitation. We further evaluated the OFIC approach with even lower input samples, then used this method to determine how the proteome is impacted by loss of superoxide dismutase *sod-1*, the ortholog of human *SOD-1*, a gene associated with amyotrophic lateral sclerosis (ALS). Analysis of 8,800 proteins from only 50 worms as the initial input showed that loss of *sod-1* affects the abundance of proteins required for stress response, ribosome biogenesis, and metabolism. In conclusion, our streamlined OFIC approach, which can be broadly applied to other systems, minimizes sample loss while offering the simplest workflow reported to date for *C. elegans* proteomics analysis.

## Introduction

The nematode *Caenorhabditis elegans* is one of the most widely used organisms to study fundamental biological processes and model human diseases,^1,2^ due to ease of culture and genetic manipulation, short lifespan, and high reproductive rate.^3^ Approximately 50% of the genes in the human protein-coding genome have *C. elegans* orthologs,^4^ allowing investigations in worms to shed light on conserved pathways. Direct analysis of *C. elegans* proteins in genetic mutants or strains grown under different conditions can lead to better understanding of complex biological processes. Technologies in mass spectrometry (MS)-based proteomics have advanced tremendously in the past two decades, enabling simultaneous identification and quantification of thousands of proteins in complex proteomes.^5,6^ To examine the *C. elegans* proteome, worms are typically lysed to extract proteins, then subjected to proteolytic digestion followed by liquid chromatography (LC) and tandem mass spectrometry (MS/MS) analysis.^7–10^ Since these worms have an exoskeleton, a tough cuticle largely comprised of cross-linked collagens,^11^ the animals must be flash frozen in harsh lysis buffer followed by grinding, homogenization, bead beating, and/or sonication to obtain sufficient protein yield.^12,13^ However, such physical disruption procedures not only cause significant loss of material,^9^ which is not optimal for low-input proteomics, but also introduce technical variation in sample preparation that can impact downstream data interpretation.

To combat these challenges, we recently proposed an on-filter in-cell (OFIC) processing based enhanced, efficient, effective and economical E4technology for global proteome analysis.^14^ Compared to other existing methods, it bypasses the need for cell lysis by directly digesting proteins in methanol-fixed cells. This significantly reduces the number of steps and amount of time required for sample preparation, and streamlines all proteomic processing in a single device.^14^ This technique is based on innovative application of an established method as cell fixation by methanol is a common practice in histochemical and cytochemical studies.^15,16^ Further, methanol fixation is the best preservation method for scRNA-seq analyses of neural cells,^17^ showing no bias in gene expression and providing the most similar profile to fresh cells. Methanol solubilizes lipids on the cell membrane and removes free water,^18^ allowing cells fixed by methanol to become porous and permeable to proteolytic enzymes. Among a variety of bottom-up proteomic sample preparation methods, in-cell digestion is quite new. Kelly and coworkers showed the first experimental evidence of in-cell processing, where the cells were fixed by formaldehyde followed by permeabilization with methanol and then in-cell digestion.^19^ Hatano *et al.* digested cells treated by methanol alone and obtained equivalent proteomic performance compared to SDS-based lysate, thus avoiding unwanted protein crosslinking by formaldehyde.^20^ We further examined the in-cell digestion method by performing all the steps on chromatographic filter devices, instead of in microtubes. As the reagents after each reaction on the filters can simply be depleted via centrifugation, this prevents chemical carryover that could potentially affect subsequent analysis. The functional resins in the filter also enable immediate peptide cleanup and desalting after digestion, which further reduces sample loss associated with conventional “drying-resuspension” procedures.^14^ The OFIC method also opens new windows for direct peptide enrichment on the filters right after protein digestion.

In our original study, we tested OFIC processing using yeast and mammalian cells, and demonstrated proof-of-concept evidence for ultra-fast and low-cell proteome analysis.^14^ However, the similarity between proteomes derived from OFIC digestion compared to lysate-based digestion methods was not evaluated. Further, whether OFIC digestion could be used to process other cell types was not assessed. Here, we utilize this advanced sample preparation technology for proteomic analysis of *C. elegans*, which is notoriously difficult to process. Using E3filters with 200 worms, we compare OFIC processing side-by-side with two common lysis-based methods, sodium dodecyl sulfate (SDS) and trifluoroacetic acid (TFA), and show the efficiency and effectiveness of the OFIC approach for low-input proteomics. We then utilized E4tips with an even lower worm input to define how the proteome is impacted by loss of evolutionarily conservedCu/Zn superoxide dismutase 1 (SOD-1),^21,22^ which plays a critical role in cellular defense via catalysis of superoxide radicals into less harmful oxygen and hydrogen peroxide. Gain of function mutations in *SOD1* cause ∼1 in 5 cases of familial amyotrophic lateral sclerosis (ALS), a fatal neurodegenerative disease characterized by motor neuron loss^23–26^ and current therapeutic approaches seek to reduce SOD1 to mitigate toxic effects.^27–29^ Children homozygous for loss-of-function *SOD1* mutations^30–32^ as well as *Sod1* knockout mice^33–35^ have significant motor system deficits, while loss of *sod-1* in *C. elegans* leads to degeneration of glutamatergic neurons upon exposure to oxidative stress^36^ and affects aging.^37,38^ Using OFIC processing of wild-type and *sod-1* mutant worms, we identified previously unknown changes in the proteome resulting from loss of *sod-1* that could have significant impact on cell function and potentially to shed light on mechanisms underlying these debilitating phenotypes.

## Experimental section

### *C. elegans* culture and collection

All *C. elegans* strains were maintained on nematode growth medium (NGM) with OP50 *E. coli* food source at 20°C. The wild-type worms used in the evaluation experiments (Figures 1-4) were Bristol N2. To analyze the impact of loss of *sod-1* on the proteome, *sod-1(hen25)*, a deletion allele generated by CRISPR / Cas9 genome editing, was used.

**Figure 1.**
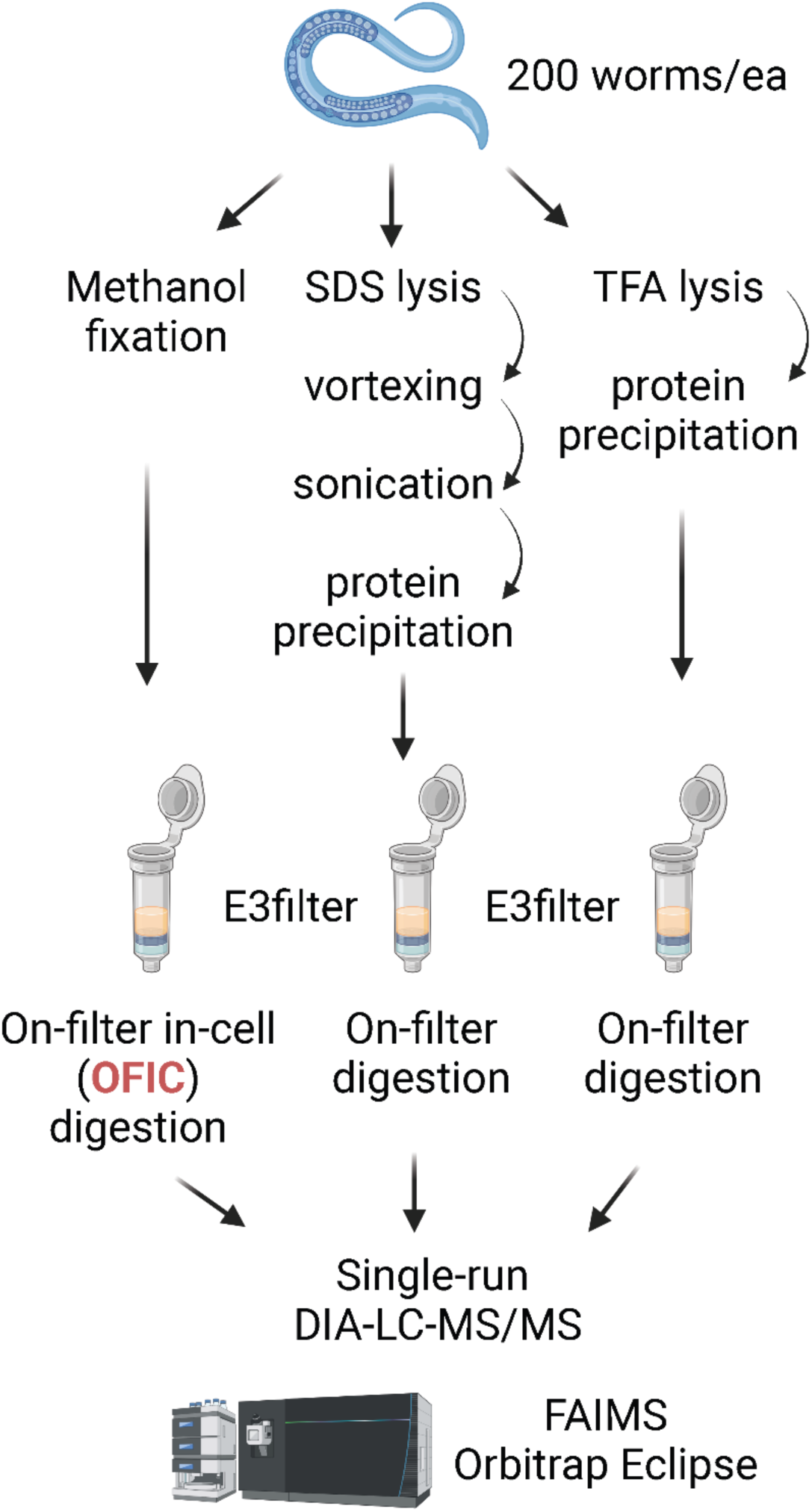
Illustrative flowchart for rapid and in-depth proteome profiling of *C. elegans*. For the OFIC processing method, the worms were fixed with methanol followed by protein digestion using E3filters. For comparison, SDS- and TFA-lysed worms were also processed on E3filters in parallel.

### Synchronization of *C. elegans*

A bleach prep using non-starved gravid adults was performed as described^39^ to obtain staged *C. elegans* for experiments shown in Figures 1-4. Isolated eggs in M9 buffer were placed on a rotator overnight at 15°C to isolate synchronized first larval (L1) stage animals. The number of L1s in 5 µl M9 buffer was counted on a microscope slide, then L1s were pipetted onto an NGM plate and grown at 20 °C until they were adults, 24 hours post fourth larval (L4). For the *sod-1* experiment, *C. elegans* were synchronized by picking late L4 stage animals, identified by vulva appearance, to a fresh NGM plate with OP50. Six hours post L4, after completion of molting, but before in-utero accumulation of eggs, the animals were briefly picked to a non-seeded NGM plate to crawl free of the *E. coli* food source. For each replicate, 50 worms total (approximately 10 at a time), were picked directly into E4tips that were pre-filled with 200 µl of pure methanol; the worm pick was visually inspected repeatedly to confirm transfer.

### OFIC processing and protein digestion of *C. elegans* with the E3filter

Live, staged *C. elegans* were rinsed three times with M9 buffer, then once with water to remove the OP50 *E. coli* food source. The worms were then processed with either on-filter in-cell (OFIC) digestion, or were first lysed with SDS buffer and TFA, before on-filter digestion as described below. For OFIC processing, a previous described protocol was followed with minor modifications.^14^ Briefly, 200 μl of pure methanol was added to aliquots of worms (three biological replicates of 200 worms), and incubated on ice for 30 minutes. The fixed worms were transferred to E3filters (CDS Analytical, Oxford, PA), and spun at 1,500 x g for one minute. The filters were washed one time with 200 μl of methanol. Then 200 μl of 50 mM triethylammonium bicarbonate (TEAB) with 10 mM Tris(2-carboxyethyl) phosphine (TCEP) and 40 mM chloroacetamide (CAA) was added, followed by incubation at 45⁰C for 10-15 min. The E3filters were then washed one time with 200 μl 50 mM TEAB. For protein digestion, 200 μl 50 mM TEAB plus 0.8 μg of trypsin was added to the samples and incubated at 37⁰C for 16-18 hours with gentle shaking (350 rpm/min). After digestion, the samples were acidified with 1% formic acid (final concentration) and left on the bench for a few minutes. In the meantime, C18 spin columns (CDS Analytical, Oxford, PA) were activated with 200 μl of methanol and spun at 1,500 x g for one min. The flow through was discarded, and the C18 spin columns were equilibrated with 200 μl of 0.5% acetic acid in water. The C18 spin columns were spun again and the flow through discarded. The E3filters were then stacked onto the C18 spin columns, 200 μl of wash/equilibration buffer (0.5% acetic acid in water) was added, and then spun at 500 x g for 15 min. This step was repeated one more time. The two stacked filters were transferred to new collection tubes, then two sequential elutions with Elution buffer I (60% ACN and 0.5% acetic acid in water) and Elution buffer II (80% ACN and 0.5% acetic acid in water), respectively, were performed. The elutions were pooled, then dried in the SpeedVac, and stored at -80⁰C until further analysis.

To generate SDS lysates, the aliquots of worms were mixed with 100 μl of SDS lysis buffer (4% SDS, 100 mM Tris-HCl, pH8.0), vortexed at 1,200 rpm for 5 min, and then sonicated under water bath for 10 min. The lysates were then mixed with 10 mM TCEP plus 40 mM CAA, and boiled at 95⁰C for 10 min. After cooling down, the samples were processed following the E3 procedure as reported previously.^14^ In brief, the lysates were mixed with 80% ACN to induce protein precipitation, then transferred to E3filters (CDS Analytical, Oxford, PA), and washed 2-3 times with 80% ACN; samples were spun at a speed of 1,000 x g and the flow through discarded. The samples were digested and desalted following the same procedure as described above.

For TFA lysate processing, the aliquots of worms were first pelleted to discard extra water and then incubated with 50 μl of pure TFA at room temperature for 5 min. The lysates were mixed with 4x volume of pure acetone to induce protein precipitation, and transferred to E3filters. This was followed by two washes with 200 μl of acetone and one wash with 200 μl of 50 mM TEAB. The samples were then mixed with 10 mM TCEP and 40 mM CAA in 200 μl of 50 mM TEAB, and incubated at 45⁰C for 10-15 min. The E3filters were then washed two times with 200 μl of 50 mM TEAB and the digestion and peptide desalting procedures were the same as described above.

### Digestion of proteins in wild type and *sod-1* mutant *C. elegans* with E4tips

Three biological replicates (50 worms each) of *sod-1* mutant and wild-type (WT) worms were picked, and transferred immediately to E4tips (CDS Analytical, Oxford, PA) that were pre-filled with 200 μl of pure methanol. The tips were incubated at 4°C for 15 min, then spun at 2,000 x g for one minute. The tips were washed one time with 200ul of methanol, and then incubated with 10 mM TCEP and 40 mM CAA (final concentrations) in 50 μl of 50 mM TEAB solution at 45⁰C for 10-15 min. The tips were spun to discard flow through and washed one more time with 200 μl of 50 mM TEAB solution. For digestion, 0.4 μg of trypsin in 100 μl of 50 mM TEAB solution was added to each sample, and then the samples were incubated at 37⁰C for 16-18 hours with gentle shaking (350 rpm/min). After digestion, the samples were acidified with 1% formic acid and spun at 500 x g for 10-15 minutes, followed by a wash step with 200 μl of wash/equilibration buffer. The E4tips were then transferred to new collection tubes, and eluted with two sequential elutions using 200 μl of Elution buffer I and Elution buffer II, respectively. The elution was pooled, dried in the SpeedVac, and stored under -80⁰C until further analysis.

### LC-MS/MS analysis

The LC-MS/MS analysis was performed on an Ultimate 3000 RSLCnano system coupled with an Orbitrap Eclipse mass spectrometer and FAIMS Pro Interface (Thermo Scientific). The peptides were resuspended into 20 μl of LC buffer A (0.1% formic acid in water), and then loaded onto a trap column (PepMap100 C18, 300 μm × 2 mm, 5 μm; Thermo Scientific) followed by separation on an analytical column (PepMap100 C18, 50 cm × 75 μm i.d., 3 μm; Thermo Scientific) flowing at 250 nL/min. A linear LC gradient was applied from 1% to 25% mobile phase B over 125 min, followed by an increase to 32% mobile phase B over 10 min. The column was washed with 80% mobile phase B for 5 min, followed by equilibration with mobile phase A for 15 min. For the ion source settings, the spray voltage was set to 1.8 kV, funnel RF level at 50%, and heated capillary temperature at 275°C. The MS data were acquired in Orbitrap at 120K resolution, followed by MS/MS acquisition in data-independent mode following a protocol described previously.^40^ The MS scan range (m/z) was set to 380-985, maximum injection time was 246ms, and normalized AGC target was 100%. For MS/MS acquisition, the isolation mode was Quadrupole, isolation window was 8 m/z, and window overlap was 1 m/z. The collision energy was 30%, Orbitrap resolution was 30K, AGC Target was 400K, and normalized AGC target was 800%. For FAIMS analysis, a 3-CV experiment (−40|-55|-75) was applied.

### Proteome quantitation and data analysis

The mass spec data were processed using Spectronaut software (version 19.1)^41^ and a library-free DIA analysis workflow with directDIA+ and the *C. elegans* protein database (UniProt 2024 release; 27,448 sequences). Briefly, the settings for Pulsar and library generation include: Trypsin/P as specific enzyme; peptide length from 7 to 52 amino acids; allowing 2 missed cleavages; toggle N-terminal M turned on; Carbamidomethyl on C as fixed modification; Oxidation on M and Acetyl at protein N-terminus as variable modifications; FDRs at PSM, peptide and protein level all set to 0.01; Quantity MS level set to MS2, and cross-run normalization turned on. Bioinformatics analyses including t-test, correlation, volcano plot and clustering analyses were performed using Perseus software (version 1.6.2.3), Prism GraphPad (version 10), and InstantClue (version 0.12.2) unless otherwise indicated. The MS raw files associated with this study have been deposited to the MassIVE server (https://massive.ucsd.edu/) with the dataset identifiers MSV000095584.

## Results and discussion

### Evaluation of OFIC processing for *C. elegans* proteome analysis

Physical disruption methods are usually employed to break *C. elegans* tissues and extract proteins for digestion and proteomics analysis. It was unknown whether a lysis-free in-cell processing method could be used to digest proteins in worms, so we first sought to assess the feasibility of OFIC for *C. elegans* proteome analysis. For comparative purpose, we processed worms in parallel using two conventional lysis methods (**Figure 1**); lysis with SDS buffer followed by vortexing and water-bath sonication, and lysis with pure TFA, which has recently shown effectiveness in processing tough samples such as gram-positive bacteria and microbiome samples.^42^ For protein digestion, E3filters were used for all comparison experiments in order to minimize technical variation.^14^ *C. elegans* proteomics experiments typically require a large input of thousands of worms.^7,10,13^ Because the OFIC processing method is naturally loss-free, we decided to use only 200 worms in our initial evaluation experiment. To increase detection sensitivity in downstream LCMS analysis, we employed high-field asymmetric-waveform ion mobility spectrometry (FAIMS), a gas-phase fractionation technique that tends to reduce the chemical background and enhance identification rate for low-input samples.^43^ Data-independent acquisition (DIA) has recently evolved as a powerful alternative to conventional DDA-based shotgun approach for highly reproducible proteome profiling.^44^ However, to our knowledge, single-shot LC-DIA MS/MS in combination with FAIMS had not been tested for *C. elegans* proteomics.

Using the OFIC processing approach with a single-shot LC-FAIMS-MS/MS run of 2.5-hour acquisition, we identified nearly 9,500 non-redundant protein groups and over 100,000 unique peptides from the 200-worm samples (Supporting Information Table S1). The identification rate was very consistent among triplicate experiments, suggesting that digesting proteins directly in methanol-fixed *C. elegans* is indeed feasible. When compared with the two lysis-based methods, the OFIC approach provided better performance in the number of protein and peptide identifications (**Figure 2 A-B**). Meanwhile, among all the protein and peptide hits identified by OFIC, about 94% and 79% of them were also identified by SDS and TFA methods, respectively (**Figure 2H, and Supporting Information Figure S1A**). This indicates that the lysis-free OFIC processing is as powerful as the lysis-based proteomics methods. For OFIC-derived proteins, the median of coefficient of variation (CV) was 6.4%, better than SDS (8.6%) and TFA (10.3%) based methods. On the peptide level, OFIC also showed the least variation among the three methods, 13.5% versus 17.1% for SDS and 19.6% for TFA (**Figure 2 C-D**). We attribute these improvements to the lysis-free and one-device nature of the OFIC method, which not only reduces sample loss but also minimizes variations during sample handling. Quantitatively, the intensity distribution of the proteomes derived from the three processing methods was largely similar (**Figure 2F**). Unsupervised clustering analysis showed some distinctness of the TFA-based processing method while clustering of OFIC and SDS methods was tighter (**Figure 2I**). Overall, the proteome-level reproducibility of OFIC digestion was high (Pearson > 0.98), and outperformed the other two lysis-based methods (**Figure 2G**).

**Figure 2.**
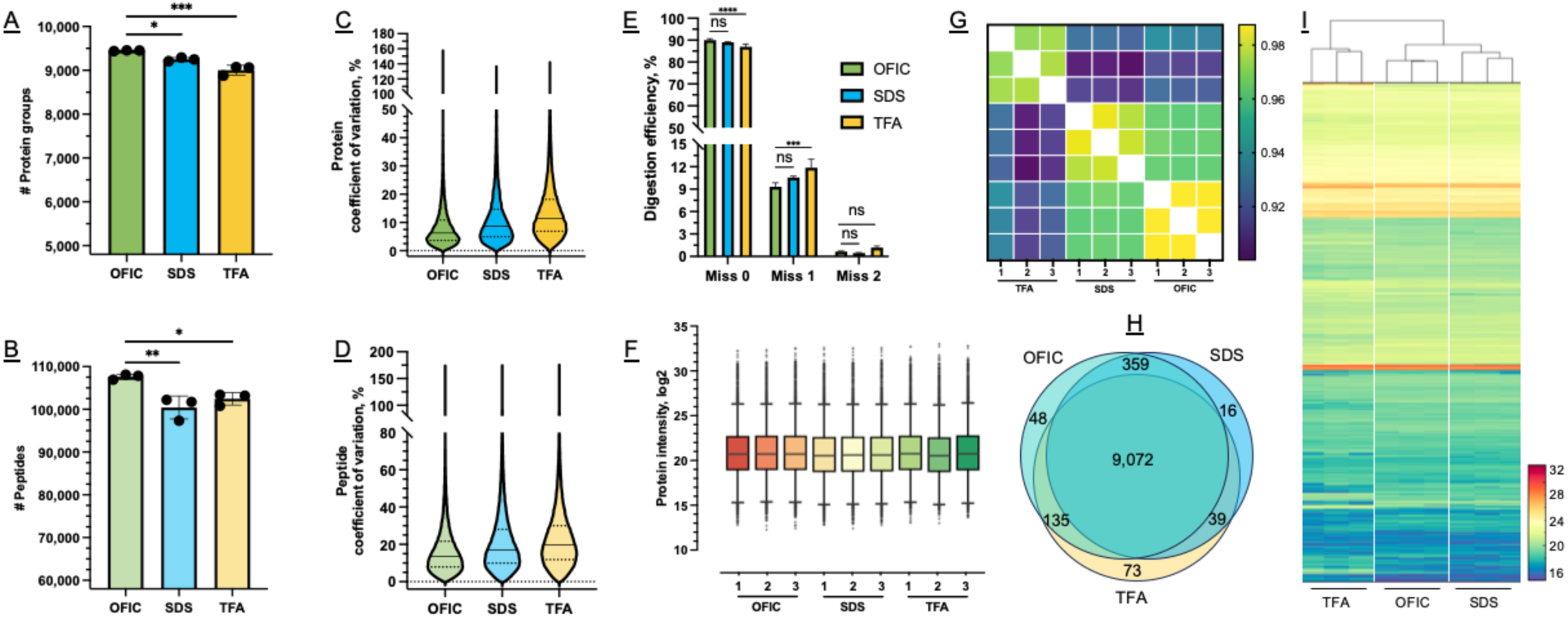
Evaluation of OFIC processing for *C. elegans* proteome analysis. (<u>A</u>-<u>B</u>) Comparison of the number of proteins and peptides derived from the three processing methods. Three replicate experiments were performed for each method. (<u>C</u>-<u>D</u>) Coefficient of variation analysis for protein and peptide identifications. (<u>E</u>) Digestion efficiency calculated based on peptide intensity. (<u>F</u>) Box plot of protein intensity. (<u>G</u>) Pearson correlation of replicate experiments between all three methods. (<u>H</u>) Venn diagram of protein identifications. (I) Heatmap of all the proteins quantified by the three methods.

We next assessed the digestion efficiency of the in-cell processing method. By peptide counts, around 23% and 3% of the peptides derived from OFIC digestion carried one and two missed cleavage sites, respectively (**Supporting Information Figure S1B**). This distribution is not significantly different from the two lysis-based processing methods assessed, and similar to recent tip-based^45^ and filter-based digestion of cell lines and muscle tissue samples.^46,47^ By peptide intensities, these miscleaved peptides contributed to less than 10% of the overall intensity of the OFIC-derived peptides (**Figure 2E**), suggesting a high quantitative yield of fully cleaved peptides by in-cell digestion.^50,51^ Taken together, we conclude that direct in-cell digestion of proteins in worms is feasible and offers distinct advantages for global analysis of the *C. elegans* proteome.

Methionine oxidation, which typically occurs during the sample preparation of bottom-up proteomic experiments,^48,49^ can hinder protein digestion and electrospray ionization, complicating the investigation of this biologically important modification that occurs *in vivo*.^50^ Therefore, we next examined the percentage of method-induced methionine oxidation. Our data showed that about 10% of methionine-containing peptides were oxidized, consistent with previous on-filter digestion experiments,^48^ and the OFIC method produced the least summed intensity of oxidized peptides (**Figure 3A**), only 0.5% or less of the overall precursor intensity. By contrast, the two lysis-based methods exhibited at least twice as much methionine oxidation. This data again highlights the in-cell process method, which not only simplifies sample handling, but also minimizes artificial modifications.

**Figure 3.**
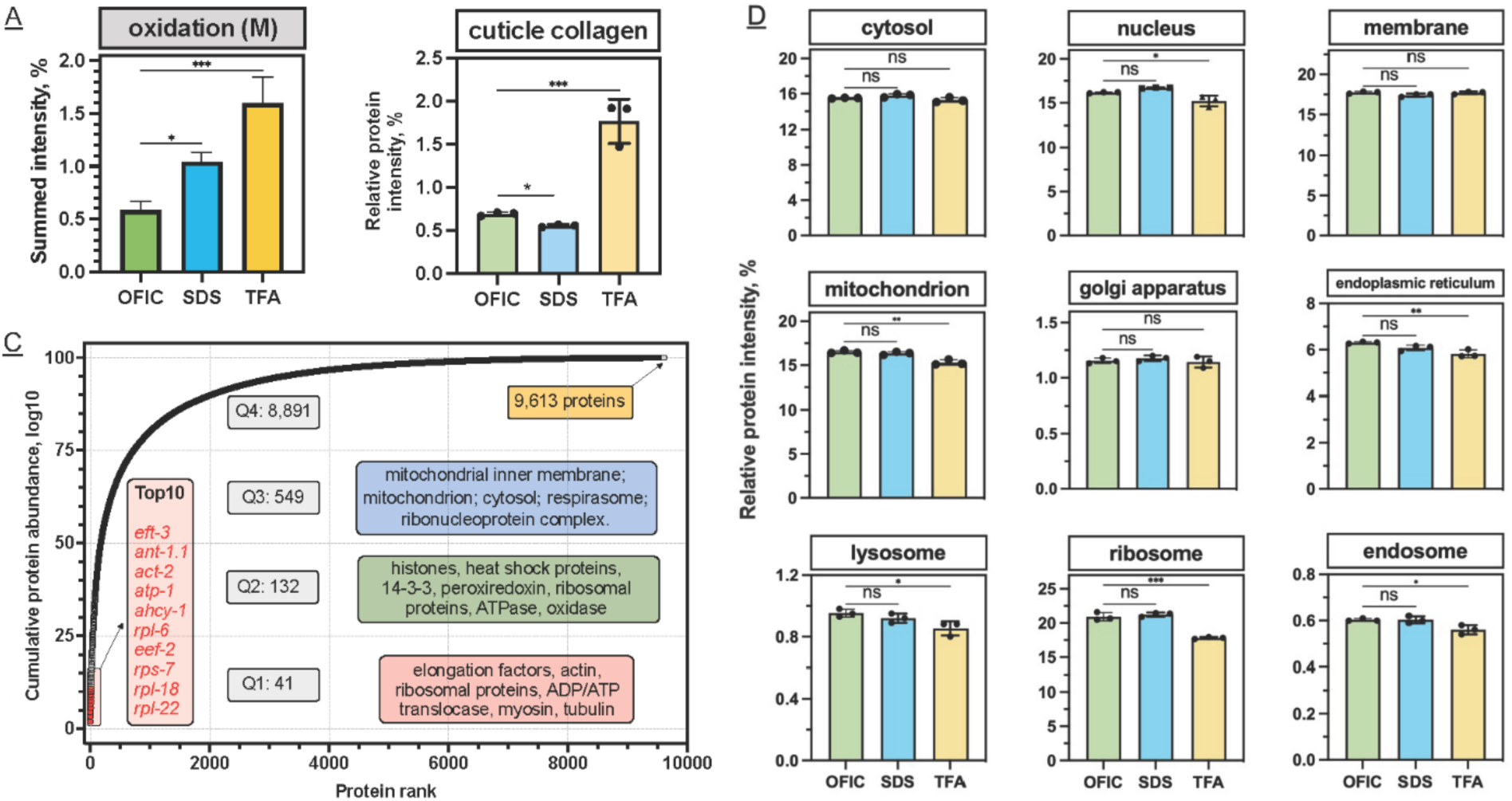
Dissecting the *C. elegans* proteome identified using the OFIC approach. (<u>A</u>) Intensity distribution of methionine oxidation among the three processing methods. The sum intensity of oxidized methionine containing peptides was divided by the total intensity of precursors for each run. (<u>B</u>) Distribution of collagen proteins among the three methods. The raw sum intensity of cuticle collagen proteins was divided by the total intensity of all identified proteins for each run. (<u>C</u>) Total quantified *C. elegans* proteins identified by OFIC processing, ranked by abundance (iBAQ intensity). The top 10 most abundant proteins, the number of proteins in each quartile and the functional classifications are as indicated. (<u>D</u>) Comparison of the intensity distribution of nine different groups of proteins based on subcellular localization for the three different processing methods. Error bars indicate triplicate experiments in panels A and D.

Next, we took a closer look at the overall *C. elegans* proteome obtained by OFIC processing, totaling 9,613 proteins after taking the mean quantity of the three biological replicates (**Supporting Information Table S1**). Notably, an in-depth study which used thousands of worms in combination with extensive peptide-level fractionation and a total of 320 hours of LCMS acquisition time identified 9,398 proteins,^7^ suggesting that our single-shot LCMS analysis of OFIC processed samples, even without any pre-fractionation, was capable of similar in-depth proteome profiling. The OFIC-derived *C. elegans* proteome spanned over seven orders of magnitude (**Supporting Information Figure S1C**), and was dominated by a small proportion of proteins. The 41 most abundant proteins comprised 25% of the total proteome mass, while the most abundant 173 proteins comprised half of the total protein mass (**Figure 3C**). These most abundant proteins include small and large ribosomal subunit proteins (e.g., RPS-7, RPS-22, RPL-6, RPL-18), cytoskeleton proteins (e.g., actin ACT-2, myosin UNC-54, tropomyosin LEV-11, and tubulin TBA-2, TBB-2) as well as nuclear proteins such as histones.

We then determined if there was a quantitative bias toward any particular functional or subcellular protein groups. The proteomes of OFIC-processed *C. elegans* and SDS worm lysates were largely not different in the identification of cytosol, nuclear, membrane, mitochondrion, Golgi apparatus, or ER proteins (**Figure 3D**). As a particular example, collagens are components of the *C. elegans* cuticle, the exoskeleton of the free-living worms that serves as a barrier between the animal and its environment. Previously, in-cell digestion was found to be ineffective for collagen proteins in mouse liver.^20^ Here, among the 173 predicted cuticle collagens.^11^ We successfully identified nearly 100 of these proteins using OFIC digestion, even more than detected in the SDS lysates (**Figure 3B**). It is likely that some cuticular collagens were not identified because our sample consisted of only adult worms and different collagen proteins cycle at specific developmental stages.^11^ TFA is a strong acid substitute of detergents for biological sample lysis, and can often dissolve cells and tissues completely.^42^ Our data showed significantly more collagen proteins from TFA lysate (**Figure 3B**), suggesting that TFA may be a good alternative to collagen proteomics studies, at the expense of relatively large technical variations and under-representation of certain protein classes such as nuclear, mitochondrion, and ribosome (Figure 3D). In conclusion, methanol permeabilization of the *C. elegans* cuticle permits enzyme accessibility to intra-organismal regions, enabling direct digestion of proteins in fixed worms and can be used as an efficient approach for processing intact *C. elegans* for proteome analysis.

### Applying the OFIC method to low-input *C. elegans* samples

Having established that OFIC processing of 200 worms achieved similar proteome depth to approaches with larger input and extensive fractionation, we set out to investigate if we could further reduce the initial input for in-cell processing. In this experiment, we utilized the tip-based E4tip method, a smaller version of the E4 filter devices, to further reduce non-specific sample loss on the plastic surface.^49^ With E4tips and single-run LC-FAIMS-DIA MS/MS, we were able to identify on average 6,055 proteins from five worms (**Figure 4**). As the input increased from 5 to 10, 20 and 40 worms, we consistently gained more identifications, though we note that dilutions from an aliquot of worms was used, and thus these numbers are approximations. From 50 worms, we were able to identify 8,857 proteins and 107,826 peptides. As the input increased to 200 worms, the gain in peptide level was less than 1%, although the number of proteins increased by nearly 7%. These data show the feasibility of in-cell processing for ultra-low worm proteome analysis and suggest that the single-run LCMS used in our study may have reached near maximal capacity with a sample of 200 worms.

**Figure 4.**
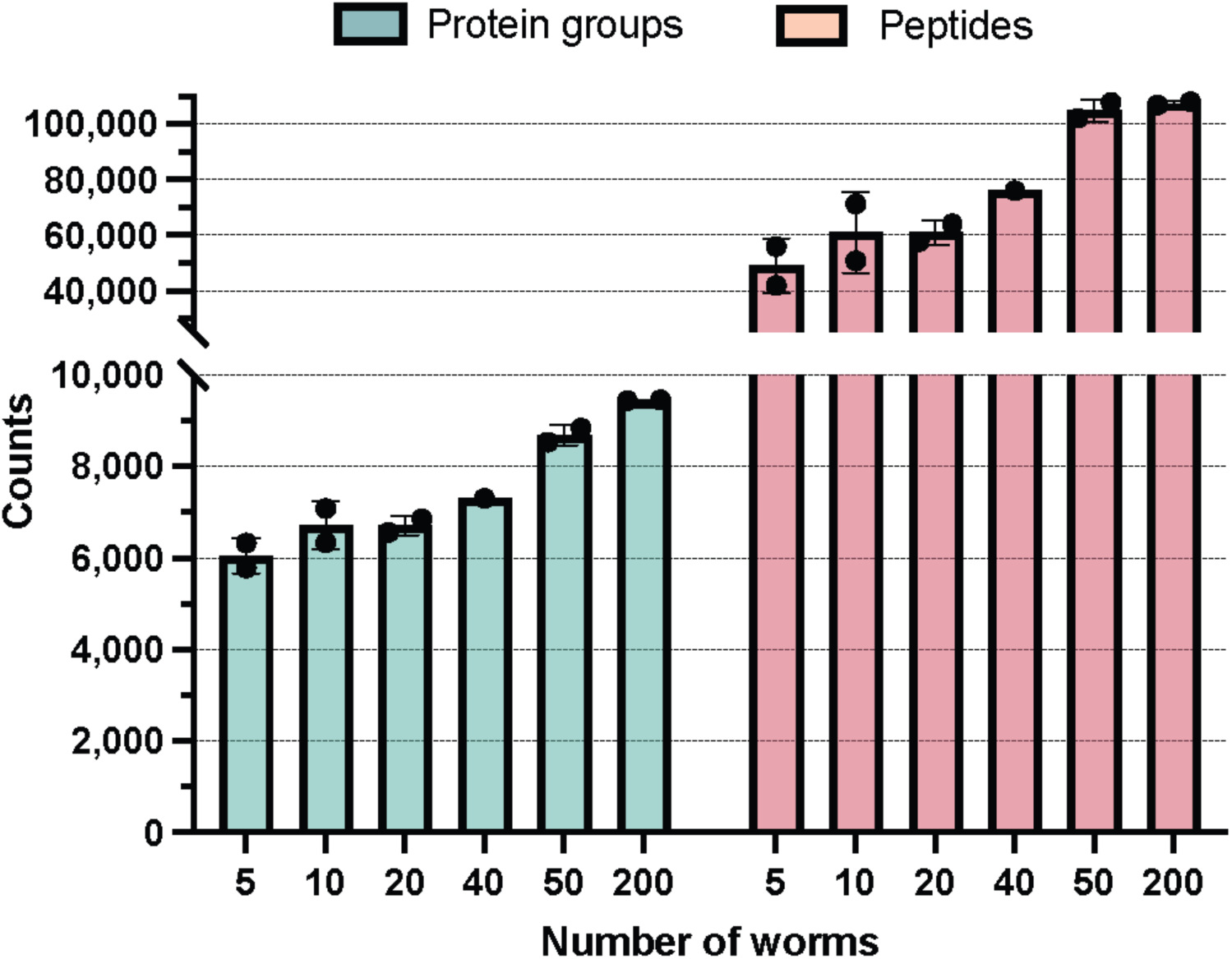
Applying OFIC digestion to study the *C. elegans* proteome with ultra-low input. Protein (left) and peptide (right) identifications summarized for different input of worms. Error bar indicates duplicate experiments.

### Loss of SOD-1 impacts the global *C. elegans* proteome

One of the best features of the fully integrated sample preparation with E4tips is that the proteomic work flow is extremely simple and nearly loss-free. Worms can be picked off plates and loaded directly into the tip, allowing the animals to be fixed immediately. This prevents perturbations that could alter the physiology of the animals, thus making the quantitative proteomics data more biologically meaningful.^44^ Having shown the power and usefulness of the OFIC digestion method for low-input intact *C. elegans* proteomics sample processing, we sought to use this method to compare the proteome of wild-type and mutant *C. elegans* (**Figure 5A**). Autosomal dominant mutations in *SOD1* cause ALS through a gain of function mechanism^25,26^ and a recently approved therapeutic approach aims to reduce *SOD1* expression^27^. However, since complete loss of SOD1 function causes debilitating motor system abnormalities^30–32^, it is important to understand the consequences of lowering this gene product. Thus, we sought to use the OFIC approach developed in this study to determine how loss of *C. elegans sod-1*, which leads to an increase in both cytosolic and mitochondrial superoxide levels^51^, impacts the global proteome. To do so, we utilized E4tips to analyze 50 worms per sample, performing three biological replicates for both WT and *sod-1* mutant strains. On average, we quantified 8,812 proteins from the six experiments (**Supporting Information Table S2**). Compared to the wild type control, the abundance of 332 proteins in the *sod-1* mutant exhibited a ≥1.5-fold change, with significance defined as an adjusted q value < 0.05 (**Figure 5B**). We then utilized WormCat, an online tool for analysis of *C. elegans* datasets^52^ to categorize the 155 upregulated and 177 downregulated proteins (**Figure 5 C,D**).

**Figure 5.**
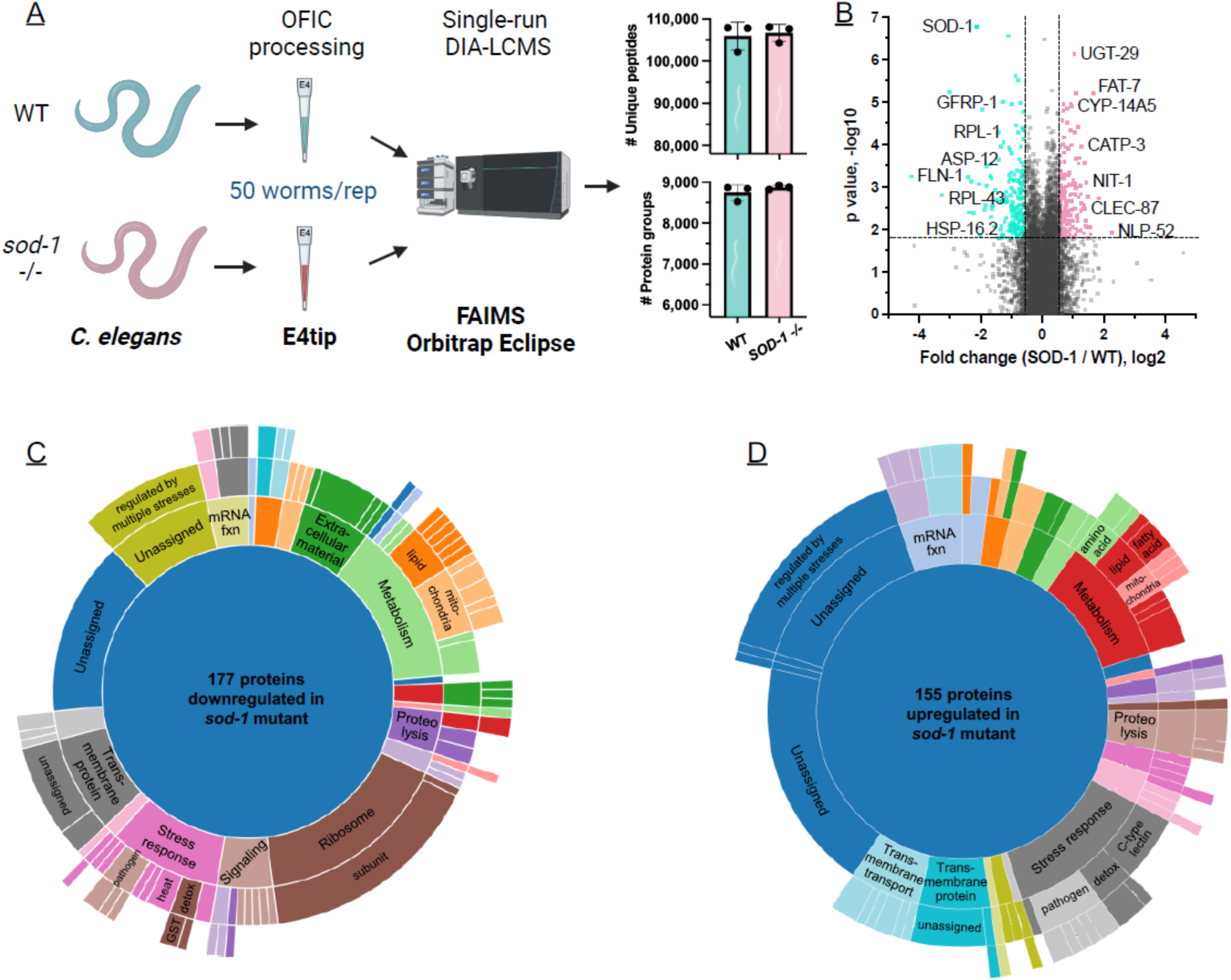
Use of OFIC processing to investigate the impact of SOD-1 on the proteome. (<u>A</u>) Overview diagram of the experimental workflow. (<u>B</u>) Volcano plot shows proteins with decreased (blue) and increased (red) abundance in the *sod-1* mutant; dotted lines show the 1.5-fold change and and adjusted q value 0.05. (<u>C</u>)(<u>D</u>) Sunburst plots display relative abundance of up- and down-regulated proteins in each category; concentric rings provide additional information about specific categories.

Consistent with redox dysregulation, there was a significant increase in stress response proteins in the *sod-1* mutant, including glutathione S-transferase GST-24, and UDP glycosyltransferases UGT-26, UGT-29, and UGT-31. Notably, other stress response proteins, including heat shock proteins HSP-16.1 and HSP-16.2 and glutathione S-transferases GST-29 and GST-39 were significantly downregulated, suggesting that loss of *sod-1* causes global changes in stress response (**Supporting Information Table S3**). Abundance of certain proteins involved in lipid metabolism, including the acyl-CoA synthetases ACS-1 and ACS-2, as well as the stearoyl-CoA desaturase FAT-7, was increased in the *sod-1* mutant. Several proteins involved in fatty acid metabolism are differentially regulated in ALS model mice with a SOD1(G93A) mutation,^53^ which causes decreased enzyme activity,^54^ suggesting that the impact of SOD-1 on metabolism is conserved. Examination of the proteins with reduced abundance in the *sod-1* mutant showed a highly significant enrichment of ribosomal proteins (p < 10^-20^) including 19 RPL proteins, which are in the large 60S ribosomal subunit (**Supporting Information Table S3**). Previously, knockout of *SOD1* was shown to affect biogenesis of the 60S ribosomal subunit in KRAS mutant lung cancer cells^55^ and our data show that loss of SOD-1 has potential to impact ribosome abundance in non-cancerous cells in vivo. These results also suggest that loss of *sod-1* may cause reduced translation in older animals, which could have a substantial impact on cell function.

## Conclusions

Here we have presented an in-cell processing method for *C. elegans* proteome analysis using low-input samples. Unlike typical bottom-up proteomics which require long processing procedures, this innovative single-vessel approach is extremely convenient as it bypasses cell lysis and disruption, requires no further sample transferring after loading, and digests proteins directly in the cells, which substantially reduces sample loss and technical variation. We systematically compared OFIC digestion with on-filter digestion of samples lysed with SDS or TFA and showed that OFIC digestion does not impact the number of protein and peptide identifications, reproducibility, digestion efficiency, undesired modification, and identification of proteins localized to subcellular compartments. Further, OFIC digestion is superior to conventional lysis-based processing methods in terms of simplicity and accessibility and could easily be scaled up for high-throughput experiments using our recently developed E4 filter devices,^14^ which enable sample cleanup with functional resins to provide LCMS-ready samples.

Single vessel methods, pipette tip-based vessels in particular, are of particular interest to the proteomics community due to their small reaction volume, reduced nonspecific adsorption to surfaces, and great ease of handling, scaling up, and automation.^45,56^ In existing methods, digestion-compatible or LCMS-friendly reagents are used to lyse cells and perform digestion, however, these chemicals may lead to a biased proteome or affect LCMS analysis.^57,58^ In our study, we have convincingly demonstrated that OFIC treatment can not only substantially simplify proteomic sample processing, but also enable unbiased deep proteome profiling of samples with low input. The proteomic comparison of wild-type and *sod-1* mutant worms using OFIC-based E4tips showed the great potential of this simple method for answering biologically significant questions.^57^ We discovered that loss of *sod-1* impacts not only the abundance of proteins involved in stress response, but also metabolic and ribosomal proteins. These previously unknown changes in the proteome resulting from *sod-1* depletion could have significant impact on cell function and SOD1-related conditions. A potential drawback of this study is that we did not examine the OFIC method for single worm proteomics due to the lack of access to top-of-the-line LC and MS instruments. However, we hypothesize that OFIC sample processing using E4tips could greatly facilitate ultralow-cell and single-cell proteomics. For instance, E4tips could possibly be connected directly to a downstream data acquisition system such as Evosep LC, thus making it even simpler than the recently reported One-Tip method.^45^ We anticipate that the OFIC approach will greatly benefit *C. elegans* proteomics community, and can be applied to a wide range of organisms and cell types beyond.

## Acknowledgements

We would like to acknowledge support from the National Institute of General Medical Sciences (NIGMS) under award number P20GM104316 for the Orbitrap Eclipse MS instrument. This work was supported by NIH-NIGMS T32 GM133395 (to M.E. as part of the Chemistry Biology Interface predoctoral training program), and NIH-NIGMS R01 GM135433 (to J.E.T.). Special thanks to the R&D team of CDS Analytical for fruitful discussions of E4filter devices. The table of content figure was created with BioRender.

---

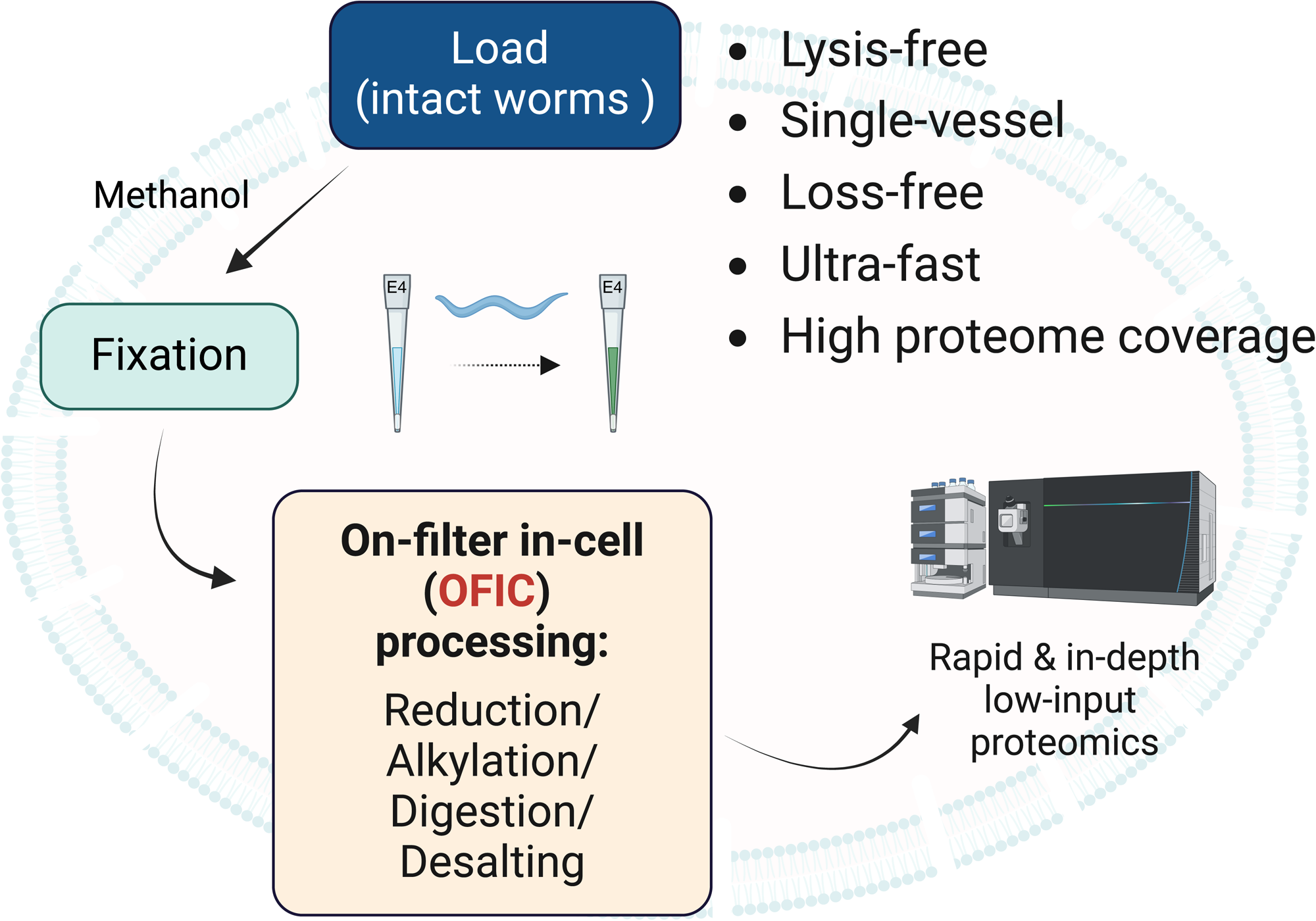

