## Supplemental Figure 1 and methods for "In-cell processing enables rapid and in-depth proteome analysis of low-input *Caenorhabditis elegans*"

##### Supporting information available

**Supplementary Figure S1**. Analysis of *C. elegans* proteome derived from OFIC processing.

**Supplementary method**. Detailed experimental protocol and flowchart of OFIC method.

**Supplementary Table S1**. Detailed list of proteins identified by the OFIC, SDS and TFA methods (Excel).

**Supplementary Table S2**. Detailed list of proteins associated with *sod-1* mutant and wild-type *C. elegans* strains (Excel).

**Supplementary Table S3.** Detailed WormCat analysis of proteins with altered abundance in the *sod-1* mutant strain (Excel).

**Supplementary Figure 1**. Analysis of the *C. elegans* proteome derived from OFIC processing. (A) Venn diagram shows the overlaps of the unique peptide sequences derived from the three methods. (B) Digestion efficiency assessed by peptide counts. (C) The dynamic range of the *C. elegans* proteome derived from the OFIC method.

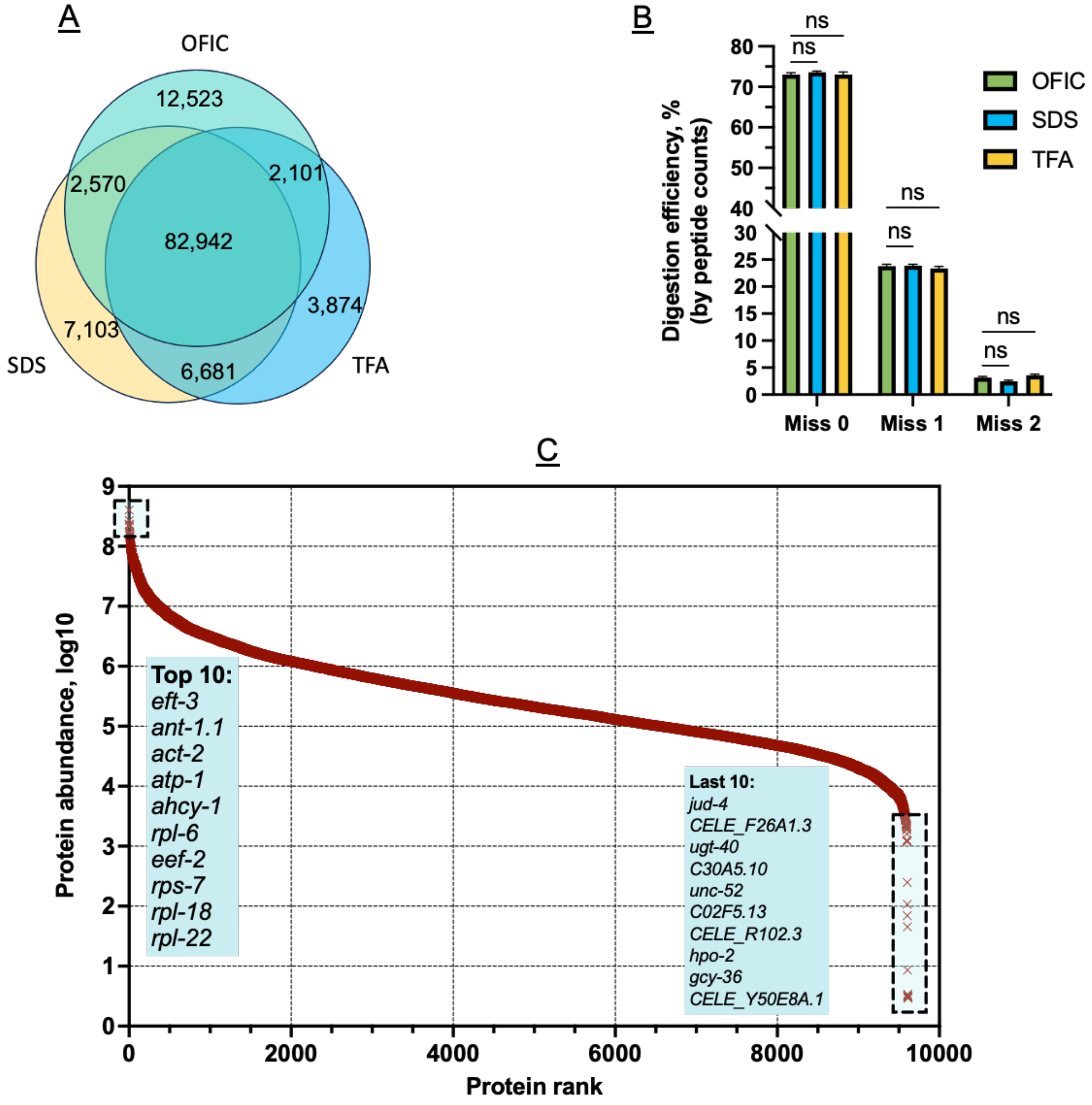

### On filter in-cell digestion (OFIC) of fresh *C. elegans* worms using E4technology

1. **Worm treatment**
   Rinse live *C. elegans* three times with M9 buffer, then one time with water, to remove *E. coli* food source.
2. **Sample loading**
   Pick worm manually (approximately 5-10 at a time), and transfer directly into E4filters
   that are pre-filled with 200 μl of pure methanol. Visually inspect worm pick to confirm transfer.
   Note: Estimated capacity for E4tip, < 500 worms, and E4 spin columns, 200-2,000 worms.

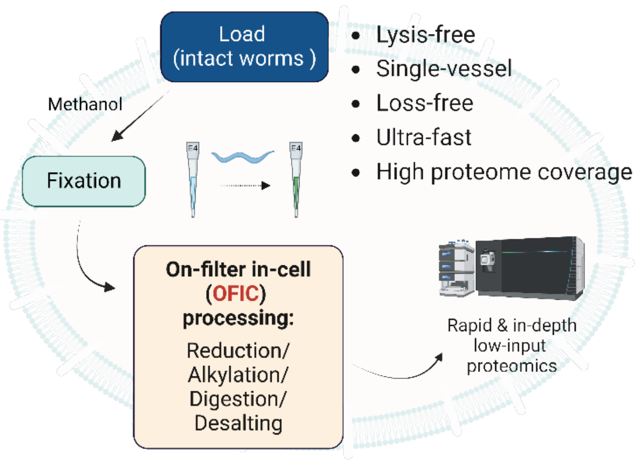


3. ***C. elegans* fixation**
   Incubate E4filters at room temperature for 15 min. Centrifuge at 1,500 x g for two min, discard flow through. Add 200 μl of methanol, repeat this step one more time.
   Note: the flow through may be collected here for metabolomics analysis.
4. **Reduction and alkylation**
   Add 100 μl of 50 mM triethylammonium bicarbonate (TEAB) with 10 mM Tris(2-carboxyethyl)phosphine (TCEP) and 40mM chloroacetamide (CAA); incubate at 45°C for 10 min with gentle shaking.
5. **Wash**
   Add 200 μl of 50 mM TEAB solution, centrifuge at 1,500 x g for two min, discard flow through.
6. **Digestion**
   Add 100 μl 50 mM TEAB, desired enzyme (Trypsin or Trypsin/Lys-C mix) at 1:50 ratio. Incubate at 37°C for 16-18 hours with gentle shaking.
   Note: no cap is required for E4tips.
7. **Acidification and desalting**
   After digestion, add formic acid to final concentration of 1%, centrifuge at 500 x g for 10 min. Add 200 μl 0.5% acetic acid in water, centrifuge at 1,500 x g for 2 min, discard flow through.
   Note: here, E4tips can be transferred to Evosep LC for direct LCMS acquisition.
8. **Elution**
   Transfer E4filters to clean collection tubes, do two sequential elution by adding 200 μl 60% acetonitrile/0.5% acetic acid in water (elution I), and 80% acetonitrile/0.5% acetic acid in water (elution II), centrifuge at 1,500 x g for 2 min to collect elution to the same tube. Dry samples in the SpeedVac, and store at -80°C. The peptides are now desalted and ready for LCMS analysis.
